# Remdesivir but not famotidine inhibits SARS-CoV-2 replication in human pluripotent stem cell-derived intestinal organoids

**DOI:** 10.1101/2020.06.10.144816

**Authors:** Jana Krüger, Rüdiger Groß, Carina Conzelmann, Janis A. Müller, Lennart Koepke, Konstantin M. J. Sparrer, Desiree Schütz, Thomas Seufferlein, Thomas F.E. Barth, Steffen Stenger, Sandra Heller, Alexander Kleger, Jan Münch

**Affiliations:** Department of Internal Medicine 1, Ulm University Hospital, Albert-Einstein-Allee 23, Ulm, Germany; Institute of Molecular Virology, Ulm University Medical Centre, Meyerhofstrasse 1, Ulm, Germany; Department of Pathology, Ulm University Hospital, Albert-Einstein-Allee 23, Ulm, Germany; Institute for Microbiology and Hygiene, Ulm University Medical Center, Ulm 89081, Germany

**Author notes:** **Corresponding authors**: Prof. Dr. Jan Münch, Institute of Molecular Virology, Ulm University Medical Centre, Meyerhofstrasse 1, 89081 Ulm, Germany, Prof. Dr. Alexander Kleger, Department of Internal Medicine I, Ulm University, Albert-Einstein-Allee 23, 89081 Ulm, Germany. contributed equally (Jana Krüger, Rüdiger Groß, Carina Conzelmann, Janis A. Müller, Sandra Heller, Alexander Kleger, Jan Münch). Author contributions J.K., R.G., C.C. and J.A.M. acquired, analyzed and interpreted data, drafted and revised the work. J.K. performed stem cell differentiation, organoid culture, immunohistochemistry stainings and fluorescent microscopy. R.G., C.C. and J.A.M. performed and analyzed infection experiments with organoids as well as Caco-2 cells and performed qPCR and viability assays. L.K. and K.S. performed confocal imaging of stained organoids, deconvolution and editing of microscopy pictures and revised the work. D.S. established work with EK1. T.F.E.B. provided sections of gastrointestinal organs for immunofluorescence stainings and helped with the analysis. S.S. supervised BSL3 work. T.S. helped to interpret data. S.H. A.K. and J.M. directed the work, interpreted the data and drafted the manuscript with input from all authors.

## Abstract

Gastrointestinal symptoms in COVID-19 are associated with prolonged symptoms and increased severity. We employed human intestinal organoids derived from pluripotent stem cells (PSC-HIOs) to analyze SARS-CoV-2 pathogenesis and to validate efficacy of specific drugs in the gut. Certain, but not all cell types in PSC-HIOs express SARS-CoV-2 entry factors ACE2 and TMPRSS2, rendering them susceptible to SARS-CoV-2 infection. Remdesivir, a promising drug to treat COVID-19, effectively suppressed SARS-CoV-2 infection of PSC-HIOs. In contrast, the histamine-2-blocker famotidine showed no effect. Thus, PSC-HIOs provide an interesting platform to study SARS-CoV-2 infection and to identify or validate drugs.

## Introduction

The severe acute respiratory syndrome coronavirus 2 (SARS-CoV-2) infects the respiratory tract with mostly mild symptoms, but up to 20% of patients develop severe pneumonia eventually followed by multi-organ failure and death^1^. Intriguingly, up to 50% of patients present with gastrointestinal symptoms, associated with prolonged disease duration and increased severity^2^. Viral RNA is detected in rectal swabs long after nasopharyngeal swabs tested negative^3^. Infection of host cells with SARS-CoV-2 requires Transmembrane Serine Protease 2 (TMPRSS2) and Angiotensin-Converting Enzyme 2 (ACE2). Both proteins apparently mediate multiorgan tropism, as they are detected in esophagus, ileum, and colon^4^. Single layered human intestinal organoids (HIOs) derived from human gut express ACE2 and are susceptible to SARS-CoV-2^5^. However, HIOs are less complex in architecture and lack *in vivo* transplantability, in contrast to human intestinal organoids derived from pluripotent stem cells (PSC-HIO). Albeit the exploration of new drugs is rapidly evolving, knowledge on their efficiency to inhibit intestinal infection of SARS-CoV-2 is unknown. However, drug testing might require a more complex organotypic culture system to reflect the true value of a given drug. Remdesivir is up to now the sole agent showing benefit on pulmonary phenotypes in COVID-19 patients^6^. The histamine-2-blocker famotidine has been suggested to reduce severe COVID-19 course^7^, but experimental or clinical proof is lacking. Here, we employ PSC-HIOs to study SARS-CoV-2 tropism with respect to distinct intestinal cell types and test drug candidates in a human organotypic culture system resembling natural 3D environment.

## Methods

Detailed information about stem cell culture, intestinal differentiation, virus strains, infection of organoids, drug testing, RT-qPCR and histology of organoids and tissue sections is presented in the supplemental text.

## Results

### ACE2 and TMPRSS2 are expressed in the gastrointestinal (GI) tract and in PSC-HIOs

First, we studied expression of ACE2 and TMPRSS2 in organs of the GI. Expression of both SARS-CoV-2 entry factors was most prominent in the epithelial lining of duodenum (**Fig.1A**), gallbladder and colon as compared to Caco-2 cells (**Fig.S1A,B**). Duodenal cells show apical ACE2 expression in the glycocalyx and a strong cytoplasmic TMPRSS2 signal. Cells of gallbladder and colon show a strong luminal expression of both proteins while sections of gastric mucosa and esophagus only reveal weak expression. These findings suggest that various cell types in gastrointestinal tissues are SARS-CoV-2 target cells, though with different expression intensity. Next, we analyzed ACE2 and TMPRSS2 expression in *in vitro* differentiated PSC-HIOs (**Fig.S1C**). Both proteins were readily expressed in organoids (**Fig.1B;Fig.S1D,E)** mimicking expression patterns in human tissue (**Fig.S1A,B**). Moreover, ACE2 is expressed on chromogranin A (CHGA)-positive enteroendocrine and lysozyme (LYZ)-positive Paneth cells, while mucin 2 (MUC2)-positive goblet cells appeared weak (**Fig.1C;Fig.S1E**).

**Figure 1:**
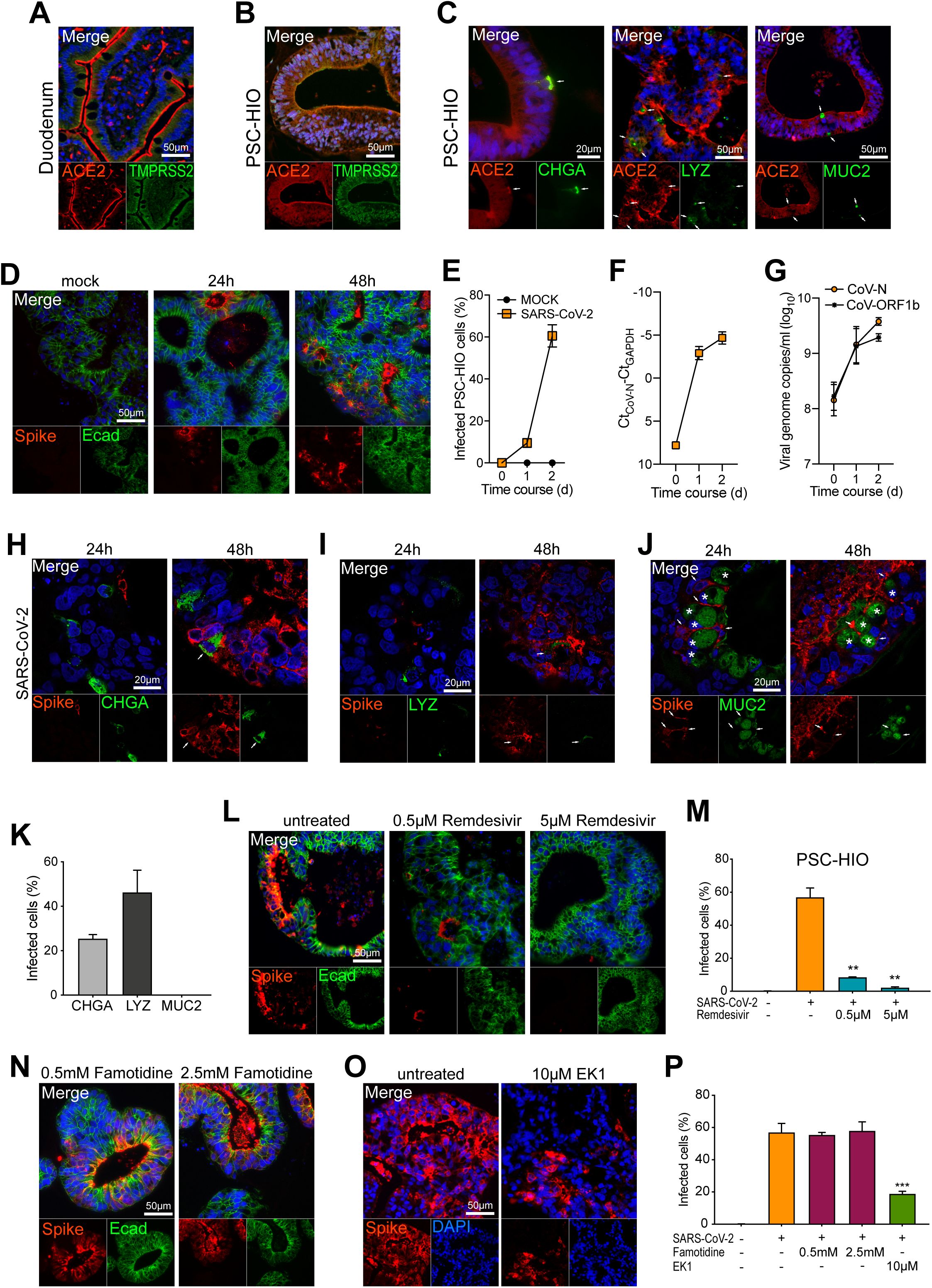
Human intestinal organoids derived from pluripotent stem cells (PSC-HIOs) express ACE2 and TMPRSS2 and support SARS-CoV-2 infection which is inhibited by remdesivir. **A-C** Duodenum biopsy and PSC-HIOs express ACE2 and TMPRSS2 (A,B). In PSC-HIOs, ACE2 is coexpressed with enteroendocrine marker chromogranin A (CHGA), Paneth cell marker lysozyme (LYZ) and goblet cell marker mucin 2 (MUC2). Nuclei are stained with DAPI in blue. Arrows indicate coexpression (C). **D** PSC-HIOs were infected with SARS-CoV-2 and stained for viral spike protein and E cadherin (Ecad) at 24 h and 48 h post infection. Nuclei are stained with DAPI in blue. **E** Infected cells of PSC-HIOs are quantified according to presence of viral spike protein compared to total cell number. **F** Relative SARS-CoV-2 viral RNA abundance in PSC-HIOs at different time points was quantified using primers amplifying CoV-N in RT-qPCR. For normalization, Ct values obtained for amplification of GAPDH from the same cultures were subtracted from the Ct value obtained for CoV-N. **G** Viral genome copy number in extracellular matrix (Matrigel) of infected PSC-HIO was assessed by RT-qPCR targeting to CoV-N and -ORF1b-nsp14. **H-K** Infected PSC-HIOs were stained for viral spike protein and chromogranin A (CHGA) at 48 h post infection. Arrow indicates coexpression (H). Co-staining was also performed with lysozyme (LYZ) as Paneth cell marker (arrow indicates coexpression) (I) and mucin 2 (MUC2) as goblet cell marker (asterisks indicate spike negative goblet cells and arrows indicate surrounding spike-positive cells) (J). Confocal imaging was performed to analyze coexpression; nuclei are stained with DAPI in blue. Specific cell types infected with SARS-CoV-2 were quantified according to presence of viral spike protein (mean ± SEM) (K). **L,M** Remdesivir treatment decreases SARS-CoV-2 infection of PSC-HIOs as shown by viral Spike protein staining 48 h post infection. Quantification of infected, spike protein positive cells (mean ± SEM). **N-P** Famotidine does not affect SARS-CoV-2 infection of PSC-HIO cells, as shown by staining of spike protein 48 h post infection (N). The presence of 10 µM EK1 during initial infection decreases the percentage of infected PSC-HIO cells as shown by viral spike protein staining 48 h post infection (O). Quantification of infected, spike protein positive cells (mean ± SEM) (P).

### PSC-HIOs support SARS-CoV-2 infection and replication

Upon SARS-CoV-2 infection of PSC-HIOs, viral spike protein was detected in 10% of the cells after 24h, which increased to 57% at 48h, suggesting viral replication and spreading infection (**Fig.1D,E**). In line, RT-qPCR revealed an increase in SARS-CoV-2 RNA levels over time in both organoids and the surrounding extracellular matrix (Matrigel) (**Fig.1F,G**). Co-staining of viral spike protein with markers for distinct intestinal cell types showed co-expression in enteroendocrine (CHGA+) and Paneth cells (LYZ+) (**Fig.1H,I,K;Fig.S1H**). In contrast, goblet cells (MUC2+) were negative for spike, yet surrounded by spike-positive cells (**Fig.1J,K**;**Fig.S1H**), suggesting that they may not or only abortively be infected. Notably, infected cells are positive for cleaved caspase 3 (CASP3) indicating initiated apoptosis in SARS-CoV-2-positive intestinal cells (**Fig.S1F,G**). Taken together, these results show that SARS-CoV-2 productively infects most cell types in gut organoids, with the notable exception of goblet cells.

### Remdesivir but not famotidine inhibits SARS-CoV-2 in a colorectal cell line and intestinal organoids

We next analyzed the effect of remdesivir and famotidine on virus infection. We determined an IC_50_ value for remdesivir of ∼46 nM in colorectal Caco-2 cells, while viability remained normal up to inhibitory concentrations of 25 µM (**Fig.S1I,J**). Notably, remdesivir decreased infection rates of PSC-HIOs by 86% at a concentration of 500 nM and almost completely abolished infection at 5 µM (**Fig.1L,M**). In contrast, famotidine concentrations of up to 2.5 mM did not affect SARS-CoV-2 infection of Caco-2 cells **(Fig.S1K,L)** and PSC-HIOs (**Fig.1N,P**). Findings were confirmed by qPCR (CoV-N) upon prolonged remdesivir and famotidine treatment (not shown). Finally, we evaluated the antiviral activity of EK1, a recently described peptidic pan-coronavirus fusion inhibitor^8^. The presence of 10 µM EK1 during infection decreased the number of spike-positive cells 48h after infection by 38% (**Fig.1O,P**).

## Discussion

We report that *in vitro* differentiated PSC-HIOs express ACE2 and TMPRSS2 and support productive infection with SARS-CoV-2 in a cell type-specific manner launching a valuable pathomechanistical model for SARS-CoV-2 infection of the gut. Notably, remdesivir and EK1 but not famotidine block SARS-CoV-2 in PSC-HIOs indicating suitability to treat gastrointestinal COVID-19. Additionally, drugs with anti-SARS-CoV-2 activity validated in PSC-HIOs might be particularly promising to ameliorate gastrointestinal symptoms in the clinic.

A study by Lamers *et al*. recently demonstrated SARS-CoV-2 infection of enterocytes in small intestinal organoids from primary gut epithelial cells^5^. We extended this study by showing infection of enteroendocrine cells in addition to enterocytes suggesting that infection may not only disrupt absorption and transport of metabolites, but also hamper hormone secretion. Similarly, infection of Paneth cells may support further disturbance of host defense by impeding secretion of antiviral and antimicrobial proteins into the gut. As COVID-19 patients suffering from gastrointestinal symptoms show prolonged disease duration and severity of disease, these cells are particularly relevant for local immune defense and, thus, drugs that suppress intestinal SARS-CoV-2 infection are of high interest.

Remdesivir, developed to treat Ebola virus infection, provides the first antiviral attempt to treat COVID-19, albeit clinical success appears heterogeneous and benefiting subpopulations remain to be defined^6^. Similarly, effectivity of this agent across distinct organs is completely unclear. We here show that remdesivir inhibits SARS-CoV-2 infection of and replication in PSC-HIOs, highlighting its possible use for treatment of gastrointestinal infection occurring simultaneously with or independently of respiratory manifestation of COVID-19.

In contrast, famotidine, proposed to inhibit proteins essential for viral replication^7^ did not affect SARS-CoV-2 infection and spread in a human epithelial colorectal adenocarcinoma cell line and PSC-HIOs. However, our current treatment regime does not allow to determine long-term effects, which may play a role in clinical deterioration. Therefore, different treatment regimens in our model systems are necessary to account for various modes of action of tested compounds. Still, retrospective data need to be interpreted with caution^7^.

## Acknowledgements

The authors would like to thank Katrin Köhn, Aref Saed, Daniela Krnavek, Nicola Schrott and Juliane Nell for their excellent technical assistance and Kanishka Tiwary and Karolin Walther for providing resources. We also thank Markus Breunig, Michael Melzer, Jessica Merkle and Meike Hohwieler for helpful discussions. R.G., C.C., L.K., D.S. and J.K. are part of and R.G. is funded by a scholarship from the International Graduate School in Molecular Medicine Ulm. This work was supported by the EU’s Horizon 2020 research and innovation programme (Fight-nCoV, 101003555 to J.M.) and by the DFG (CRC1279 to S.S. and J.M.).

## References

1. Zhu N, et al. New England Journal of Medicine 2020.

2. Wei X-S, et al. Clinical Gastroenterology and Hepatology 2020.

3. Xu Y, et al. Nature medicine 2020;26:502–505.

4. Xiao F, et al. Gastroenterology 2020;158:1831–1833 e3.

5. Lamers MM, et al. Science 2020.

6. Beigel JH, et al. N Engl J Med 2020.

7. Freedberg DE, et al. Gastroenterology 2020.

8. Xia S, et al. Science advances 2019;5:eaav4580.

