## Supplement for "Remdesivir but not famotidine inhibits SARS-CoV-2 replication in human pluripotent stem cell-derived intestinal organoids"

**Content:**

***Supplementary Figure 1***	*p. 2-3*

***Supplementary Materials and Methods***	*p. 4-6*

***Supplementary References***	*p. 6*

**Supplementary Figure 1**

**Figure S1:** **ACE2 and TMPRSS2 expression in gastrointestinal tissues and intestinal organoids, their susceptibility to SARS-CoV-2 infection and effect of remdesivir and famotidine on virus infection.**

**A,B** Paraffin embedded sections of indicated gastrointestinal tissues and embedded Caco-2 cells were stained for ACE2 and TMPRSS2 protein. Strong luminal expression is found in epithelia of duodenum, colon and gallbladder, weaker expression in the stomach and the mucosal lining of the esophagus. Representative images are shown (A). Quantitative expression of proteins was scored according to immunofluorescence stainings, using Caco-2 cells as a reference for strong expression (+++) (B).

**C,D** Intestinal organoids (PSC-HIOs) were differentiated from human embryonic stem cells and maintained in 3D Matrigel culture (C). PSC-HIO sections were stained for ACE2 and TMPRSS2 protein showing robust expression (D).

**E** Confocal imaging shows weak expression of ACE2 in MUC2-positive goblet cells. Arrows indicate coexpression.

**F,G** SARS-CoV-2 infected cells show expression of apoptosis marker cleaved caspase 3 (CASP3) 48 h post infection (F). Coexpression was confirmed by confocal imaging, arrows indicate coexpression (G).

**H** Infected PSC-HIOs were stained for viral spike protein and chromogranin A (CHGA) as marker for enteroendocrine cells at 48 h post infection. Co-staining was also performed with mucin 2 (MUC2) as goblet cell marker and lysozyme (LYZ) as Paneth cell marker. Nuclei are stained with DAPI in blue. Arrows indicate coexpression of spike with CHGA and LYZ or spike-negative MUC2-positive cells, respectively.

**I** Immuno-detection of SARS-CoV-2 in infected Caco-2 cells treated with different concentrations of remdesivir at 48 h post infection (mean ± SD). Calculated half maximal inhibitory concentration (IC_50_) is indicated.

**J** Effect of indicated concentrations of remdesivir on cell viability in Caco-2 cells (mean ± SD).

**K** Immuno-detection of SARS-CoV-2 in infected Caco-2 cells treated with different concentrations of famotidine 2 days post infection (mean ± SD).

**L** Effect of indicated concentrations of famotidine on cell viability in Caco-2 cells (mean ± SD).

**Supplementary Materials and Methods**

**Drugs:** Remdesivir was obtained from Selleck Chemicals (#S8932), famotidine from Sigma Aldrich (#F6889-1G), EK1 synthesized by the Core Facility Functional Peptidomics, Ulm University Medical Center.

**Cell culture.** Caco-2 cells were grown in Dulbecco’s modified Eagle’s medium (DMEM) supplemented with 10% FCS, 100 units/ml penicillin, 100 µg/ml streptomycin (Sigma), 2 mM L-glutamine, 1 mM sodium pyruvate, and NEAA (Sigma #M7145). Vero E6 cells were grown in the same medium but with 2.5% FCS. Cells were grown at 37°C in a 5% CO_2_ humidified incubator.

**Stem cell culture and intestinal differentiation.** Human embryonic stem cell (hESC) line HUES8 (Harvard University) was used with permission from the Robert Koch Institute according to the “156. Genehmigung nach dem Stammzellgesetz, AZ: 3.04.02/0156”. Cells were cultured on hESC Matrigel (Corning) in mTeSR1 medium (Stemcell Technologies) at 5% CO_2_, 5% O_2_, and 37°C. Medium was changed daily and cells were splitted with TrypLE Express (Invitrogen). For differentiation, 300,000 cells per well were seeded in 24-well-plates coated with growth factor reduced (GFR) Matrigel (Corning) in mTeSR1 with 10 µM Y-27632 (Stemcell technologies). The next day, differentiation was started at 80-90% confluency according to the protocol published by Hohwieler *et al.^1^.*

**Virus strains and virus propagation.** Viral isolate BetaCoV/Netherlands/01/NL/2020 (#010V-03903) was obtained from the European Virus Archive global and propagated on Vero E6 cells. Infectious virus titer was determined as plaque forming units (PFU).

***In vitro* infection and cell-based SARS-CoV-2 immunodetection assay.** To determine SARS-CoV-2 infection, 30,000 Caco-2 cells were seeded in 96-well plates. The next day, the compound of interest was added and the cells were infected with a multiplicity of infection (MOI) of 0.001 in a total volume of 180 µl. Two days later, cells were fixed in 4% PFA, permeabilized using 0.1% Triton, and stained with 1:5,000 diluted anti-SARS-CoV-2 Spike protein antibody 1A9 (Biozol GTX-GTX632604) in antibody buffer (10% FCS and 0.3% Tween 20 in PBS) for 1 h at 37°C. After 3 washes, the secondary HRP-conjugated antibody (Thermo Fisher #A16066) (1:15,000) was incubated for 1 h at 37°C. Following 4 times of washing, the TMB peroxidase substrate (Medac #52-00-04) was added for 5 min and the reaction stopped using 0.5 M H_2_SO_4_. The optical density (OD) was recorded at 450 nm - 620 nm using the Asys Expert 96 UV microplate reader (Biochrom). Values were corrected for the background signal derived from uninfected cells and untreated controls were set to 100% infection.

***In vitro* cell viability assay.** The effect of analyzed compounds on the metabolic activity of Caco-2 cells was analyzed using the CellTiter-Glo® Luminescent Cell Viability Assay (Promega #G7571) as previously described^2^.

**Infection of organoids and drug testing**. To prepare *in vitro* differentiated organoids for infection, matrigel was dissolved in Collagenase/Dispase (Roche) for 2 h at 37°C and stopped by cold neutralisation solution (DMEM, 1% BSA and 1% penicillin-streptomycin). Organoids were transferred into 1.5 ml tubes and infected in 350 µl virus inoculum containing 5 × 10^4^ PFU SARS-CoV-2 with or without drugs for 1 h at 37°C. Organoids were then resuspended in 35 µl cold GFR Matrigel to generate cell-matrigel domes in 48 well plates. After 10 min at 37°C, intestinal growth medium (DMEM F12 (Gibco), 1x B27 supplement (Fisher Scientific), 2 mM L-glutamine, 1% Penicillin/streptomycin, 40 mM HEPES (Sigma), 3 µM CHIR99021, 200 nM LDN-193189 (Sigma), 100 ng/ml hEGF (Novoprotein) and 10 µM Y-27632 (Stemcell Technologies)) was added and organoids were incubated at 37°C.

**Isolation of RNA from matrigel and organoids for RT-qPCR.** Viral RNA from matrigel of infected cells or organoids was isolated using the Qiagen Viral RNA Mini Kit (Qiagen #52906) or Qiagen RNeasy Plus Mini Kit (Qiagen #74136), respectively, both as described by the manufacturer. For this purpose, matrigel was digested as described above, mixed with AVL buffer and incubated at room temperature (RT) for 20 min before freezing at -20°C. Similarly, organoids were lysed in 600 μl RLT Plus buffer containing 1% 2-mercaptoethanol, vortexed for 30 s and then frozen at -20°C until analysis. Organoid lysates were homogenized using QIAshredder (Qiagen #79656).

**RT-qPCR.** RT-qPCR from matrigel was performed with primer sets targeting N (nucleoprotein) and ORF1b-nsp14 (ORF1b)^3^ as previously described^4^ using TaqMan® Fast Virus 1-Step Master Mix (Thermo Fisher, #4444436) and a StepOnePlus Real-Time PCR System (96-well format, fast mode). Synthetic SARS-CoV-2-RNA (Twist Bioscience, #102024) was used as a standard to obtain viral copy numbers. For RNA isolated from organoids, GAPDH was analyzed as an endogenous control (Applied Biosystems, #4310884E) and resulting Ct values subtracted from those obtained for CoV-N reactions for the same samples. All reactions were run in duplicates.

**Histology of organoids and tissue sections.** Sections of human gastrointestinal tissues were provided by the pathology department of Ulm University. Experiments were conducted in accordance with guidelines of the Ethics Committee of the Federal General Medical Council and approved by the Ethics Committee of the University of Ulm. For histological examination of organoids, they were fixed in 4% PFA over night at 4°C, washed with PBS, and pre-embedded in 2% agarose (Sigma) in PBS. After serial dehydration, intestinal organoids were embedded in paraffin, sectioned at 4 µm, deparaffinized, rehydrated and subjected to heat mediated antigen retrieval in tris Buffer (pH 9) or citrate buffer (pH 6). Tissue was permeabilized with 0.5% Triton-X for 30 min at RT, stained over night with primary antibodies in antibody diluent (Zytomed) in a wet chamber at 4°C. After washing with PBS-T, slides were incubated with secondary antibodies (Alexa Fluor IgG H+L, Invitrogen, 1:500) and 500 ng/ml DAPI in Antibody Diluent for 90 min in a wet chamber at RT. After washing with PBS-T and water, slides were mounted with Fluoromount-G (Southern Biotech). Negative controls were performed using IgG controls or irrelevant polyclonal serum (anti-mycobacterium tuberculosis) for polyclonal antibodies, respectively. Absence of background staining confirmed specificity of the primary antibodies. Images were acquired with Keyence BZ 9000 microscope, quantified with ImageJ. Laser Scanning confocal images were acquired using the Zeiss LSM710. Z-stacks were deconvoluted and edited in Huygens (Scientific Volume Imaging) and Fiji (ImageJ).

**Primary antibodies:**

| **Antigen** | **Species** | **Cat. No.** | **Company** | **Dilution** |
| --- | --- | --- | --- | --- |
| ACE2 | rabbit | ab15348 | Abcam | 1:1500 |
| ACE2 (used for costainings) | mouse | sc390851 | Santa Cruz | 1:100 |
| SARS-CoV-2 Spike | mouse | GTX632604 | Biozol | 1:500 |
| TMPRSS2 | rabbit | ab92323 | Abcam | 1:250 |
| CASP3 | rabbit | #9664 | Cell Signaling | 1:1000 |
| CHGA | rabbit | A0430 | DAKO | 1:500 |
| MUC2 | rabbit | sc-7314 | Santa Cruz | 1:200 |
| LYZ | rabbit | ab108508 | Abcam | 1:1000 |
| Ecad | rabbit | 24E10 | Cell Signaling | 1:200 |

**Supplementary References**

1.	Hohwieler M, et al. Zeitschrift für Gastroenterologie 2016;54:748-759.

2.	Muller JA, et al. Nat Commun 2018;9:2207.

3.	Chu DK, et al. Clinical chemistry 2020;66:549-555.

4.	Groß R, et al. The Lancet 2020.
